# Low hemoglobin causes hematoma expansion and poor intracerebral hemorrhage outcomes

**DOI:** 10.1101/2024.08.15.608155

**Authors:** Azzurra Cottarelli, Rayan Mamoon, Robin Ji, Eric Mao, Amelia Boehme, Aditya Kumar, Sandy Song, Valentina Allegra, Sabrina V. Sharma, Elisa Konofagou, Vadim Spektor, Jia Guo, E. Sander Connolly, Padmini Sekar, Daniel Woo, David J. Roh

## Abstract

**Objectives:** Although lower hemoglobin levels associate with worse intracerebral hemorrhage (ICH) outcomes, causal drivers for this relationship remain unclear. We investigated the hypothesis that lower hemoglobin relates to increased hematoma expansion (HE) risk and poor outcomes using human observational data and assessed causal relationships using a translational murine model of anemia and ICH.

**Methods:** ICH patients with baseline hemoglobin measurements and serial CT neuroimaging enrolled between 2010-2016 to a multicenter, prospective observational cohort study were studied. Patients with systemic evidence of coagulopathy were excluded. Separate regression models assessed relationships of baseline hemoglobin with HE (≥33% and/or ≥6mL growth) and poor long-term neurological outcomes (modified Rankin Scale 4-6) after adjusting for relevant covariates. Using a murine collagenase ICH model with serial neuroimaging in anemic vs. non-anemic C57/BL6 mice, intergroup differences in ICH lesion volume, ICH volume changes, and early mortality were assessed.

**Results:** Among 1190 ICH patients analyzed, lower baseline hemoglobin levels associated with increased odds of HE (adjusted OR per -1g/dL hemoglobin decrement: 1.10 [1.02-1.19]) and poor 3-month clinical outcomes (adjusted OR per -1g/dL hemoglobin decrement: 1.11 [1.03-1.21]). Similar relationships were seen with poor 6 and 12-month outcomes. In our animal model, anemic mice had significantly greater ICH lesion expansion, final lesion volumes, and greater mortality, as compared to non-anemic mice.

**Conclusions:** These results, in a human cohort and a mouse model, provide novel evidence suggesting that anemia has causal roles in HE and poor ICH outcomes. Additional studies are required to clarify whether correcting anemia can improve these outcomes.

## INTRODUCTION

Intracerebral hemorrhage (ICH) carries the highest morbidity and mortality of all stroke subtypes with up to 70% of ICH survivors having severe disability^1^. ICH morbidity is largely driven by ongoing bleeding, also known as hematoma expansion (HE), and hemorrhage volume. Hemorrhage volume causes direct acute brain injury in the initial hours^2–4^, as well as secondary brain injury over the ensuing hours to days, due to the neurotoxicity of hematoma breakdown products^5^. However, medical therapies targeting HE to limit hemorrhage volume and secondary brain injury processes have not yet improved ICH outcomes^6–9^. This highlights a need to investigate alternative targets that may be relevant for acute and secondary brain injury and clinical ICH outcomes.

Lower baseline hemoglobin level and anemia consistently associate with poorer ICH outcomes^10–13^. This suggests that correcting anemia may benefit these patients, particularly because correcting anemia using liberal red blood cell (RBC) transfusion strategies are actively being studied in other acute, non-ICH, brain injury settings^14–16^. However, underlying causal drivers for anemia’s relationship with poor ICH outcomes remain unclear. Beyond their role in oxygen delivery, RBCs are also critical for coagulation and hemostasis with anemic patients known to have an “anemic coagulopathy” due to impaired radial displacements of platelets and impaired platelet activation^17–22^. While small single center ICH studies identified relationships between baseline anemia, increased HE, and larger ICH volumes^10,23–25^, the generalizability of these observations is limited^26^. Most importantly, it is unknown whether anemia plays a direct causal role in these observations given the inherent limitations of human observational studies in disentangling the overlap of medical comorbidities, disease severity, and anemia. Clarifying anemia’s role in HE and ICH outcomes would set the stage for future studies aimed at correcting and/or preventing anemia to manage ICH. Thus, we investigated the hypothesis that anemia relates to increased HE risk and poor clinical outcomes in a large multi-center ICH cohort study while accounting for relevant disease severity confounders. We also used a novel translational murine model of ICH to assess whether anemia has a causal role for HE and ICH outcomes.

## METHODS

### Human ICH cohort study

Spontaneous ICH patients were enrolled between 2010-2016 into a prospective, multi-center, multi-ethnic, ERICH (Ethnic/Racial variations of IntraCerebral Hemorrhage) study. Patients with baseline hemoglobin levels and serial CT neuroimaging data were included for the current analyses. Patients with ICH etiologies secondary to trauma, vascular malformations/aneurysm, ischemic stroke with hemorrhagic transformation, and cerebral malignancy were excluded from ERICH. Additionally, patients with primary IVH, neurosurgical interventions (hemicraniectomy and/or clot evacuation) between baseline and follow-up head CT, delayed presentations (greater than 24 hours from symptom onset), and systemic coagulopathy on admission based on conventional coagulation assays (prothrombin time >20 seconds, partial thromboplastin time >50 seconds, International Normalized Ratio >1.7, platelet count < 50×10^3^ cells/uL) were excluded to minimize relevant confounders^10^.

### Hemoglobin measurements

Baseline hemoglobin levels were from clinically-drawn complete blood counts measured by the enrolling center’s clinical laboratory. Hemoglobin concentration (g/dL) was defined as a continuous variable.

### Neuroimaging outcome

Clinical CT neuroimaging scans were uploaded and analyzed for hematoma volumes using previously described semiautomated measurement approaches (Alice software; Parexel Corp, Waltham, MA)^27^. Neuroimaging acquisition was not protocolized and was obtained per clinical practice at each enrolling institution. The initial admission head CT and final follow-up head CT within 96 hours were assessed for hematoma size for HE calculations. HE was primarily defined using a well validated threshold of either ≥33% relative expansion and/or ≥6mL absolute expansion between scans^28^. HE was secondarily assessed using other validated thresholds: a) ≥33% and/or ≥6mL parenchymal expansion and/or ≥1mL IVH expansion, b) HE ≥33%^29^. Initial and final absolute hematoma volumes were assessed as continuous variables.

### Clinical outcome

Poor long-term outcomes were assessed using the modified Rankin Scale (mRS 4-6) at 3-month follow-up as the primary clinical outcome. Assessments of long-term outcomes at 6 and 12 months were also explored. Outcomes were obtained via standardized phone interviews by trained research staff^27^.

### Statistical analysis

For the human cohort study, intergroup differences between HE and non-HE groups were determined using Mann-Whitney-*U* or *t-*tests for continuous variables and χ2 or Fisher-exact tests for categorical variables. Relationships of baseline hemoglobin with HE were assessed using logistic regression models after adjusting for baseline demographics, antithrombotic medication use, baseline ICH/IVH volume, ICH location, and time to admission CT^30^. Secondary analyses were performed using alternative definitions of HE, as specified above. Linear regression models assessed relationships of baseline hemoglobin concentration with ICH volume (as a continuous log-transformed outcome variable). Sensitivity analyses were performed adjusting for kidney disease. Additional multivariable logistic regression models assessed the association of hemoglobin concentrations with poor clinical outcomes after adjusting for similar covariates and markers of ICH severity^2^. Statistical significance was judged at p-value<0.05. Analyses were performed using SPSS (IBM) and Matlab (MathWorks).

### Standard protocol approvals and patient consent

The ERICH study was approved by institutional review boards at each enrolling site. Informed consent was obtained from each patient, or surrogate, as appropriate.

#### Animal ICH study

This study was approved by the Institutional Animal Care and Use Committee (IACUC) at Columbia University Irving Medical Center. All procedures were performed using female C57/BL6 mice (Jackson Laboratory). The schematic of the experimental setup is in Figure 2.

**Figure 1:**
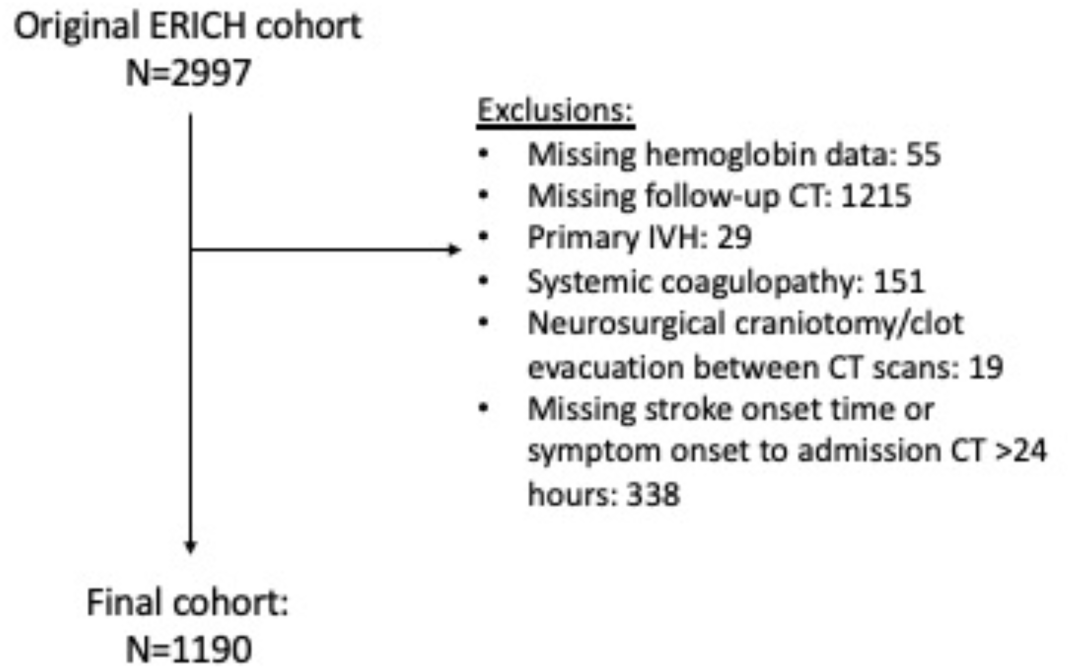
Inclusion/exclusion criteria for ERICH cohort. Legend: ERICH: Ethnic/Racial variations of IntraCerebral Hemorrhage; CT: computed tomography; IVH: intraventricular hemorrhage

**Figure 2:**
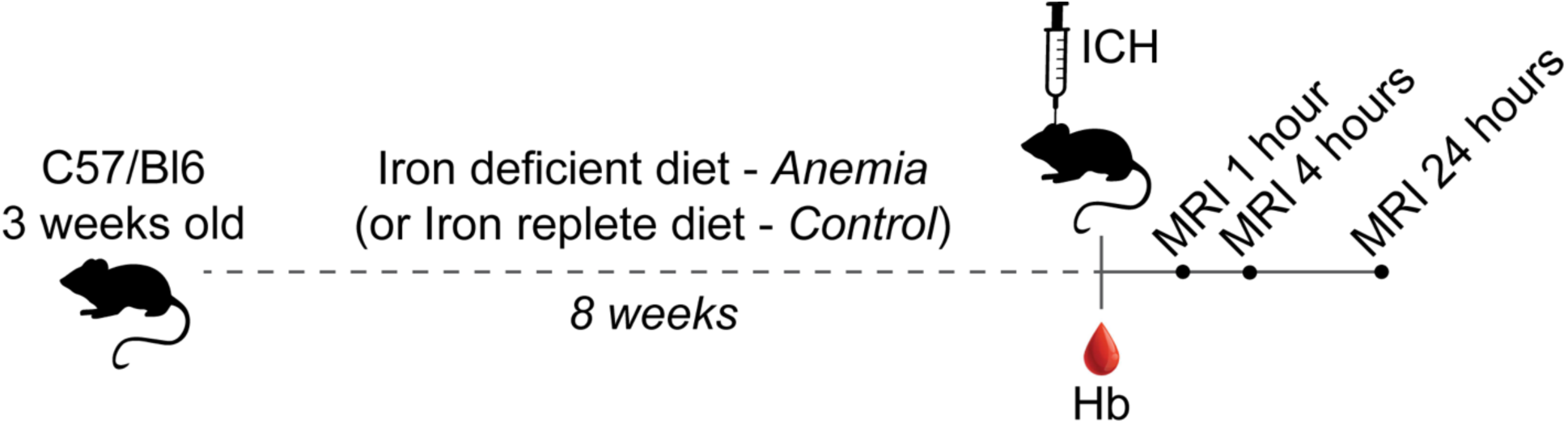
Translational murine model anemia and ICH experimental design. Legend: ICH: intracerebral hemorrhage; Hb: hemoglobin Three-week old female C57/Bl6 mice received either iron-deficient chow or iron-replete chow (for anemia vs. control, respectively) for 8 weeks. Anemia status was verified using a Drabkin assay to quantify hemoglobin levels. ICH-inducing surgery was performed in anemia vs. control mice using stereotactic injection of collagenase, followed by serial in vivo MRI neuroimaging at 1, 4, and 24 hours after ICH surgery.

### Anemia induction

Anemia was induced by feeding newly weaned, 3-week old mice for 8 weeks with either an iron-deficient diet (Envigo Teklad TD.110592) or an iron-control diet (Envigo Teklad TD.190878), the latter to provide non-anemic controls^31,32^. Successful anemia induction was verified by a quantitative colorimetric assay for determining hemoglobin concentration in whole blood using Drabkin’s reagent (Sigma). Iron deficiency was chosen as our primary anemia model as it is the underlying driver for common causes of chronic anemia (e.g., iron deficiency anemia and anemia of chronic disease)^33^. Furthermore, earlier work suggested the relevance of low iron levels in the anemia of ICH patients^34^. A separate model of anemia induction using monoclonal antibodies (i.e., anti-mouse TER-119 (BD Biosciences, 553669) and an IgG2b isotype control (BD Biosciences, 556968)) was also used to determine whether anemia *per se*, rather than iron deficiency, was the driver of neuroimaging outcomes. Given expected differences in anemia induction kinetics with anti-TER119 as compared to dietary iron deficiency, 10-12-week old female C57/Bl6 mice were injected intraperitoneally with 45μg of anti-TER119, or control, in 200μL PBS, as described^35^, 2 days before ICH induction; hemoglobin levels were quantified as described (see supplemental figure).

### ICH induction

ICH was induced in 10-12-week old female C57/Bl6 mice. Briefly, mice were anesthetized by isofluorane inhalation and their heads were immobilized in a stereotactic apparatus. After determining the injection coordinates (0.2 mm anterior, 2.3 mm lateral, 3.7 mm ventral to bregma), 0.5 U of bacterial type VII collagenase (Sigma) in 0.5 µl of PBS was injected into the right striatum at an infusion rate of 0.1 µl/min. Throughout the procedure, the animal’s body temperature was monitored and maintained at 37°C^36,37^.

### Neuroimaging/mortality outcome

Serial neuroimaging was performed at 1, 4, and 24 hours after ICH induction using a 9.4T MRI system (Bruker Biospin, Billercia, MA) and a birdcage coil (diameter: 30mm). A 3-plane scout image was used to adjust the position of the head to capture the largest area of hemorrhage. T2 weighted images (turbo RARE pulse sequence; TR/TE: 230/3.3 ms, flip angle: 70%, NEX: 4, resolution: 100um x 100um, slice thickness 400um) were then acquired^36,38^. Mice were anesthetized using isofluorane during the scan. The lesion volume was extracted and measured semiautomatically using MIPAV (Medical Imaging Processing, Analysis and Visualization software, NIH) by applying the Levelset algorithm to determine the probable boundary of the ICH lesion. Lesion HE was assessed as the primary radiographic outcome as a continuous variable (relative percent and absolute change). Initial and follow-up hemorrhage volumes (as a continuous variable) were also assessed. Early mortality data within 24 hours of ICH induction were also recorded.

### Statistical analysis

For our animal models, intergroup differences between anemic and non-anemic mice were assessed using repeated measure ANOVA, Mann-Whitney-*U* or *t-*tests for continuous variables and χ2 or Fisher-exact tests for categorical variables.

## RESULTS

### Human ICH cohort

We identified 1190 spontaneous ICH patients meeting criteria for analysis (Figure 1). The mean age was 61, 62% were male, and the cohort was racially diverse (see table 1). Baseline ICH volume was 10.6mL, baseline hemoglobin concentration was 13.8g/dL, and HE (≥33% and/or ≥6mL) was seen in 21% of patients. Intergroup differences in ICH patients with and without HE are detailed in table 1. Patients with HE were more likely to be male, with larger baseline ICH volumes, and shorter time intervals between symptom onset and admission CT. When comparing the analyzed cohort to patients excluded from analyses, there were expected differences in traditional markers of coagulopathy and symptom onset to admission CT times (i.e., greater anticoagulant medication use, INR values, and time from symptom onset to admission CT in exclusion group) given our pre-defined exclusion criteria. However, there were no differences in baseline ICH volume or ICH severity, and baseline hemoglobin levels were comparable between inclusion and exclusion cohorts (Supplemental table 1).

**Table 1:**
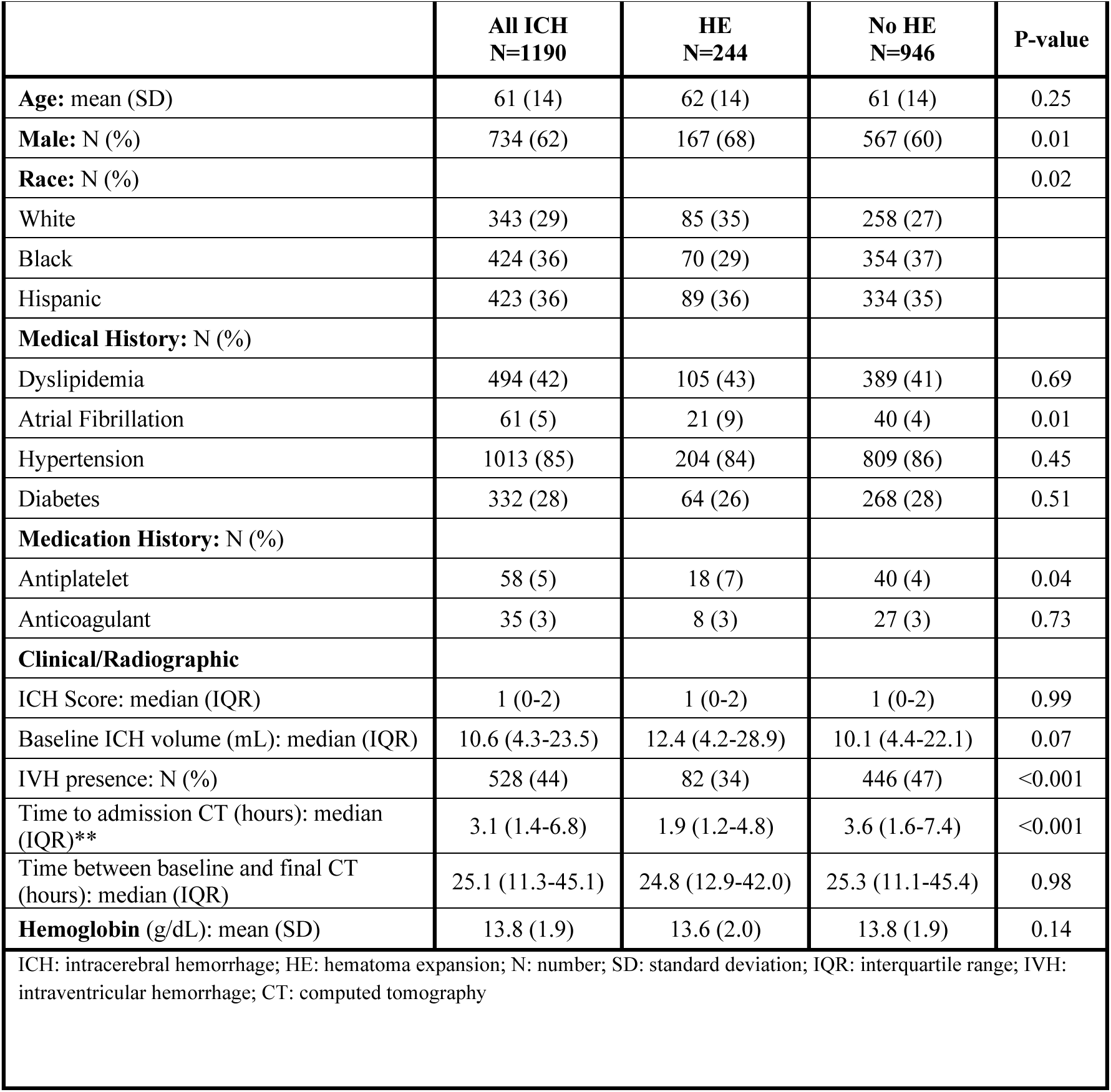
Baseline characteristics of ICH patients with and without HE.

### Relationship between hemoglobin level and HE

Using regression analyses, lower baseline hemoglobin concentration was associated with increased odds of HE (adjusted OR per -1g/dL hemoglobin decrement: 1.10; 95%CI: 1.02-1.19). Sensitivity analyses adjusting for concomitant kidney disease did not alter hemoglobin’s relationship with HE (adjusted OR per -1g/dL hemoglobin decrement: 1.09; 95%CI: 1.01-1.16). Relationships between hemoglobin level and HE were also observed when using alternative HE definitions and thresholds (table 2). No relationships between baseline hemoglobin and baseline or final ICH volume (as continuous variables) were identified using linear regression models (supplemental table 2).

**Table 2:**
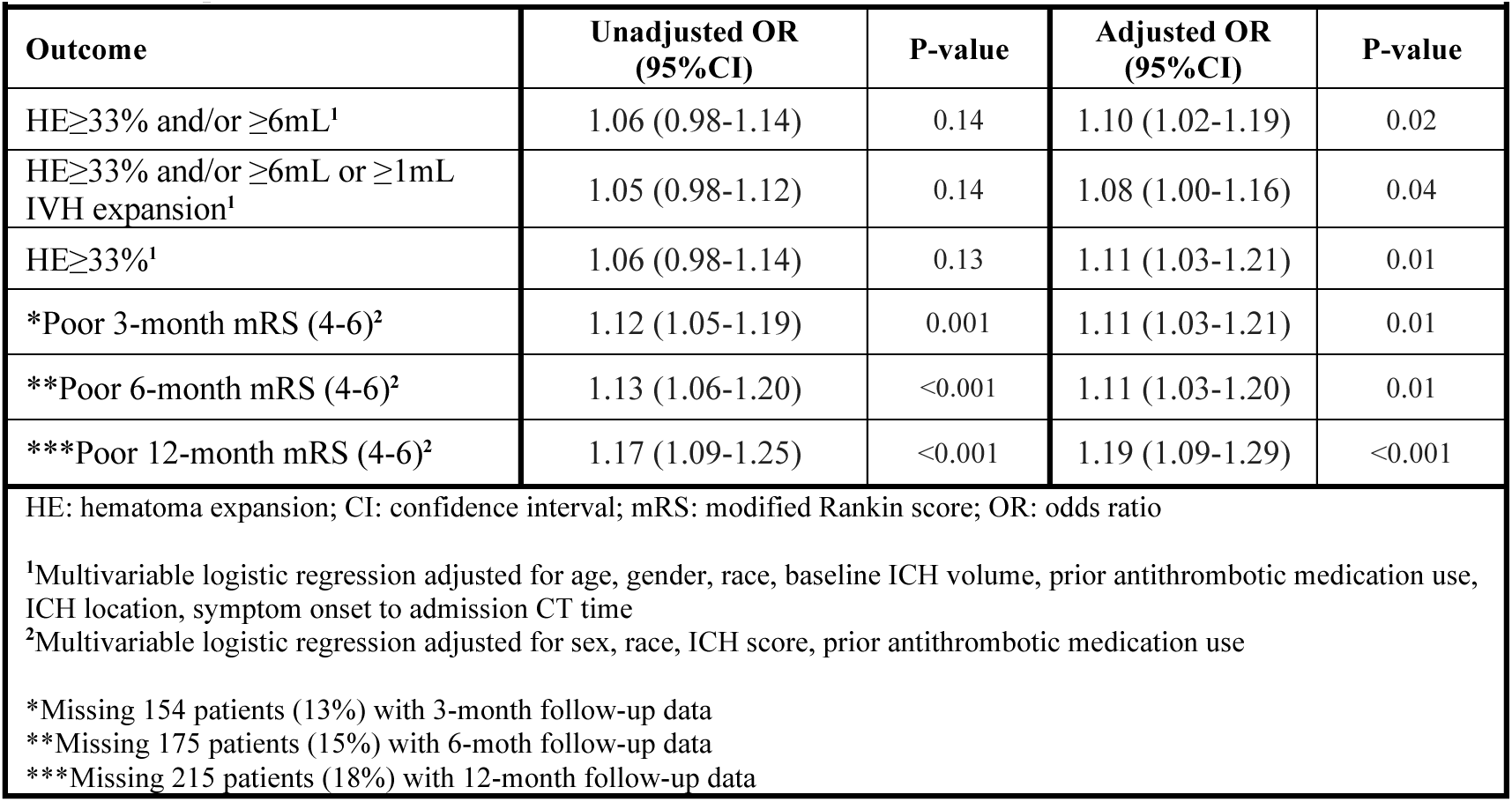
Crude and adjusted multivariable regression analysis assessing association of hemoglobin with hematoma expansion and clinical outcomes.

|  |  |  |  |  |
| --- | --- | --- | --- | --- |
| **Outcome** | **Unadjusted OR**<br>**(95%CI)** | **P-value** | **Adjusted OR**<br>**(95%CI)** | **P-value** |
| HE≥33% and/or ≥6mL<sup>1</sup> | 1.06 (0.98-1.14) | 0.14 | 1.10 (1.02-1.19) | 0.02 |
| HE≥33% and/or ≥6mL or ≥1mL IVH expansion<sup>1</sup> | 1.05 (0.98-1.12) | 0.14 | 1.08 (1.00-1.16) | 0.04 |
| HE≥33%<sup>1</sup> | 1.06 (0.98-1.14) | 0.13 | 1.11 (1.03-1.21) | 0.01 |
| *Poor 3-month mRS (4-6)<sup>2</sup> | 1.12 (1.05-1.19) | 0.001 | 1.11 (1.03-1.21) | 0.01 |
| **Poor 6-month mRS (4-6)<sup>2</sup> | 1.13 (1.06-1.20) | <0.001 | 1.11 (1.03-1.20) | 0.01 |
| ***Poor 12-month mRS (4-6)<sup>2</sup> | 1.17 (1.09-1.25) | <0.001 | 1.19 (1.09-1.29) | <0.001 |
| HE: hematoma expansion; CI: confidence interval; mRS: modified Rankin score; OR: odds ratio |  |  |  |  |
| <sup>1</sup>Multivariable logistic regression adjusted for age, gender, race, baseline ICH volume, prior antithrombotic medication use, ICH location, symptom onset to admission CT time |  |  |  |  |
| <sup>2</sup>Multivariable logistic regression adjusted for sex, race, ICH score, prior antithrombotic medication use |  |  |  |  |
| *Missing 154 patients (13%) with 3-month follow-up data |  |  |  |  |
| **Missing 175 patients (15%) with 6-month follow-up data |  |  |  |  |
| ***Missing 215 patients (18%) with 12-month follow-up data |  |  |  |  |

### Relationship between hemoglobin level and clinical outcomes

Poor 3-month outcomes were observed in 49% of the studied cohort. Lower hemoglobin levels were associated with increased odds of poor 3-month outcomes (adjusted OR per -1g/dL hemoglobin decrement: 1.11; 95%CI: 1.03-1.21). Similar relationships were observed for 6 and 12-month outcomes (table 2). Additional sensitivity analyses adjusting for kidney disease did not alter these relationships (adjusted OR per-1g/dL hemoglobin decrement: 1.11; 95%CI: 1.03-1.17).

### Relationship between hemoglobin level and ICH outcome in a murine model

Dietary iron deficiency induced significantly lower hemoglobin concentrations (6.2g/dL [+/-1.6] vs 12.4g/dL [+/-1.5], p<0.001; figure 3a). Following collagenase-induced ICH, initial baseline 1 hour MRI neuroimaging lesion volumes did not significantly differ between anemic and control mice (anemic group lesion volume mean: 3.3mm3 [+/-2.8] vs control: 2.7mm3 [+/-2.9]; p=0.95). However, anemic mice experienced significantly greater ICH lesion progression between the initial and final 24-hour MRI scans, as compared to controls (figure 3b/c). Notably, lesion expansion in anemic mice was primarily observed at later time points, between the 4- and 24-hour MRI scans, rather than between the 1- and 4-hour MRI scans (figure 3c). Additionally, final ICH lesion volumes were larger in anemic mice, as compared to controls (7.4mm3 [+/-3.4] vs 4.7mm3 [+/-4.3]; p-value=0.01) (figure 3d). Finally, mortality was more common in anemic mice, as compared to controls (figure 3e) (27% vs 0%, p-value=0.04). In experiments using an alternative anemia model utilizing anti-TER119 to induce hemolysis, significantly lower hemoglobin concentrations were observed in mice injected with anti-TER119 vs. controls (4.5g/dL [+/-2.0] vs 14.4g/dL [+/-3.1]; p<0.001). In addition, anti-TER119-treated anemic mice exhibited greater ICH lesion expansion, as compared to controls; this suggests that anemia itself, rather than underlying iron deficiency, was responsible for these findings (see supplemental figure).

**Figure 3:**
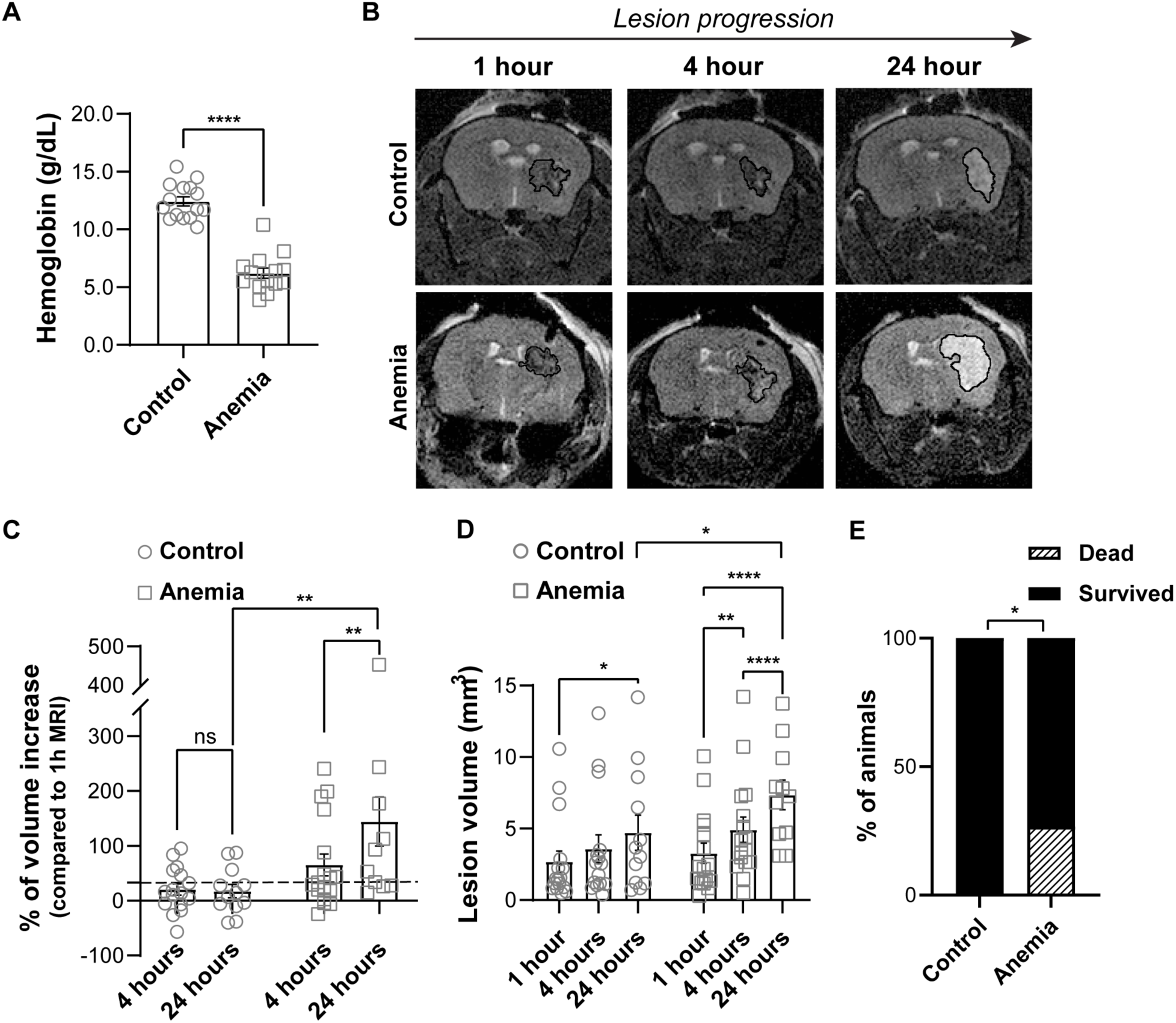
Pre-clinical murine model results. An iron-deficient chow diet results in significantly decreased hemoglobin levels, as compared to control iron-replete chow (A). Representative images of serial neuroimaging assessments of ICH lesional volume at 1, 4, and 24 hours after ICH, comparing mice with iron-deficient anemia and non-anemic controls (B). Mice with iron-deficient anemia exhibit significant increases in ICH lesion volume over time (C) and experience larger final lesion volumes, as compared to ICH control mice (D). Significantly increased mortality following ICH was seen in mice with iron-deficient anemia, as compared to controls (E). *: p<0.05; **: p<0.01; ***: p<0.005; ****: p<0.001

## DISCUSSION

While lower hemoglobin levels and anemia were consistently shown to relate to poor long-term ICH outcomes, underlying drivers for these relationships are unclear. Our translational data, leveraging a large multicenter, racially-diverse ICH patient population, paired with a novel murine model of anemia and ICH, provide critically-needed insights and investigative platforms to clarify these previous unknowns. Specifically, lower baseline hemoglobin levels associated with increased odds of HE and poor ICH outcomes in our human cohort. In addition, the murine model provides data supporting the causal role of anemia in inducing poor radiographic and clinical ICH outcomes.

In the human study, we identified relationships between lower baseline hemoglobin levels and HE across several validated definitions of HE. In addition, assessing long-term outcomes at 3, 6, and 12-months provided novel data to show that anemia’s impact on clinical outcomes is consistent even over longer follow-up time points. This is particularly important given that ICH recovery occurs over longer periods of time, as compared to other stroke types^39,40^. It was previously unknown whether baseline anemia would affect 6 and 12-month ICH outcomes. Thus, these new findings provide needed generalizability and broader insights into prior observations provided by separate smaller single center studies^10–13,23^. Collectively, these human findings suggest that the relationships between lower baseline hemoglobin levels and poor long-term outcomes are mediated, at least in part, by HE, and this effect appears to be consistent across different patient populations and racial groups. Our *a priori* exclusion of patients with systemic evidence of coagulopathy, as diagnosed by conventional coagulation assays, allowed us to assess hemoglobin as a relevant independent risk factor for HE and poor clinical outcomes, excluding confounders due to other coagulopathy etiologies or medical disease severity.

The human study was performed with the intent of disentangling comorbidities, disease severity, and anemia. Nonetheless, observational studies are inherently limited in determining causal relationships, particularly when the exposure variable (i.e., anemia) could be a surrogate assessment of the severity of the hospitalization itself and of numerous unmeasured pre-hospitalization comorbidities and hospitalization factors. Therefore, the reductionist murine model of anemia and ICH was developed to provide a preclinical model for investigating the causality of the relationships identified in human studies^10^. The collagenase model of ICH used provided an ability to assess ongoing bleeding from injured vessels captured via *in vivo* serial MRI neuroimaging. The model appeared to induce temporal ICH and HE changes in the initial hours after ICH that were translatable to the morphologic characteristics and temporal kinetics known to occur in human ICH^41^. And, the iron deficiency model of anemia was used, given that diet- or inflammation-induced iron-deficient erythropoiesis are common causes of anemia in the United States^33,42–44^, and was recently identified as a relevant driver for anemia in the setting of human ICH^34^. In this model, anemic mice experienced more HE, larger final ICH volumes, and worse clinical outcomes, as compared to non-anemic controls. Interestingly, HE in the anemic mice occurred in a more delayed fashion, primarily between the 4- and 24-hour neuroimaging scans (instead of between the 1- and 4-hour time points), as compared to non-anemic mice. Unfortunately, in the human cohort, we did not have the availability of systematic, protocolized serial neuroimaging acquisition at 3 or more time points to enable translation of these “bench findings” to the bedside. However, the human study design intentionally included final CT scan assessments at later time points, up to 96 hours after admission, to best capture any HE that could occur in a more delayed fashion beyond a hyper-acute time period. It is unclear whether the methodological approaches of prior studies, using earlier follow-up head CT scans within a 24-hour time period, prevented the identification of HE that could have occurred in anemic patients at later time points, beyond 24 hours. However, future studies will need to clarify whether anemic ICH patients do, indeed, encounter HE over longer time periods, which could have important implications for the therapeutic windows used to control hemorrhage in both research and clinical settings of HE prevention.

While these murine findings provide clarity regarding the causal role of anemia in ICH outcomes, they should be also considered as hypothesis-generating and proof-of-concept for future studies. Specifically, the precise causal mechanistic pathways by which anemia affects these outcomes should be further explored, as well as assessing whether correcting anemia will improve the prognosis. It could be speculated that these results are due to an “anemic coagulopathy” whereby radial displacement of platelets towards the vessel wall is inhibited, thereby preventing adequate platelet-endothelial interactions and clot formation^17,18,22^. This attractive hypothesis has therapeutic potential in that anemic coagulopathy can be reversed by RBC transfusion^18^. In contrast, other possible pathophysiologic mechanisms underlying the mouse model results may not be reversible by transfusion (or other therapies to correct anemia). For example, because chronic anemia affects cerebral white matter integrity^45,46^, it is possible that anemia causes HE by facilitating hematoma dispersion across impaired white matter tracts; in this case, acutely correcting anemia would be less likely to reverse these chronic pathophysiologic derangements. Furthermore, anemia may have other downstream effects, such as ischemic secondary brain injury, apart from acute HE, which may affect the nature and timing of therapeutic interventions during the clinical course. Therefore, additional murine model and human studies are required to clarify relevant biological pathways to determine which may, or may not, be acutely modifiable to improve ICH outcomes.

Despite our study’s novel investigative platform leveraging a large multicenter human cohort study with translational murine experiments to assess anemia’s causal role in ICH outcomes, there are several limitations worth noting. As previously stated, the human and murine results do not provide precise mechanistic insights. Further work is required to clarify the mechanistic underpinnings and to determine whether acute correction of anemia can improve outcomes. The animal model should also be replicated across different ages and both sexes to understand whether these factors affect ICH outcomes. Similarly, it is not yet known whether RBC transfusions in the murine model or in our human cohort will be beneficial. Although a prior human study identified a potential benefit of RBC transfusion for ICH outcomes^47^, most human observational publications concerning ICH and other disease settings suggest that RBC transfusions are associated with poor outcomes. These findings appear to be driven because RBC transfusions are often a proxy for disease severity, creating difficulties in disentangling various comorbidities and the reasons for needing an RBC transfusion from independent effects of the RBC transfusion itself^48–51^. To complicate these human observations further, there are variable efficacies and risks from each RBC transfusion itself based on individual RBC unit quality and donor characteristics^52–54^. Therefore, further pre-clinical studies will be needed to clarify whether RBC transfusions (or other interventions) can ameliorate ICH outcomes. Given the numerous etiologic drivers of anemia, there are multiple treatment options beyond RBC transfusions, including iron repletion, erythropoietin administration, and hepcidin blockade. Thus, if anemia correction is desirable, it is important to understand the underlying pathophysiology to be able to identify the appropriate therapy. Although both inflammation and iron deficiency are relevant to the anemia seen in ICH patients^34^, it is unclear whether other anemia etiologies impact the HE and ICH outcomes observed in the current study. Finally, hemoglobin levels are not static throughout a hospitalization. While we focused on baseline hemoglobin levels, given that these would precede the primary radiographic HE outcome, hemoglobin decrements and anemia development during an ICH hospitalization have been shown to be frequent, rapid, and separately related to poor outcomes. Thus, a better understanding of the drivers and outcomes of chronic anemia at presentation, and of acute anemia developing during the ICH hospital course, will be necessary to design future studies to address the potential benefits of correcting these forms of anemia.

## CONCLUSION

The results from the current human study and the pre-clinical murine model provide novel evidence to suggest that anemia has causal role in HE and poor ICH outcomes. Further work is required to clarify the mechanistic underpinnings of these findings and to understand whether correcting anemia can improve outcomes.

## Acknowledgements

Thanks to Che-Feng Chang and Lauren Sansing for their guidance and input for the collagenase ICH model leveraged for this study. In addition, thanks to Steven Spitalnik and Eldad Hod for their advice and support.

## Author contributions

DR contributed to the conception and design of the study; AC, RM, RJ, EM, AB, AK, SS, VA, SVS, EK, VS, JG, ESC, PS, DW, DR contributed to the acquisition and analysis of data; AC, RM, AB, DR contributed to drafting the text or preparing the figures.

## Availability of data

The data that support the findings of this study are available from the corresponding author upon reasonable request.

---

**Supplementary figure 1:**
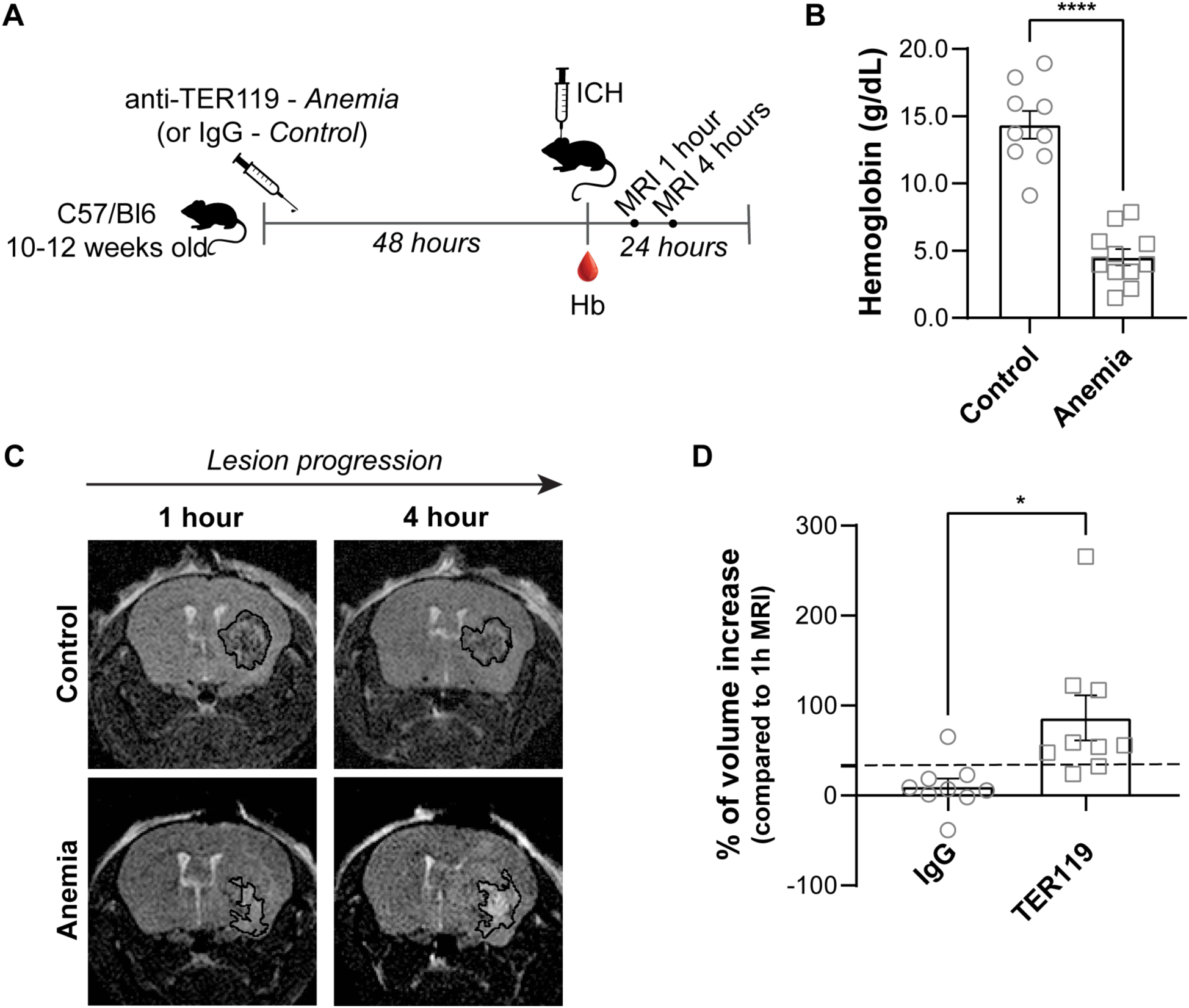
Alternative anemia ICH model using anti-TER119. Legend: ICH: intracerebral hemorrhage Experimental design for anti-TER119 experiments (A). Female 10-12-week old C57/Bl6 mice were injected intraperitoneally with anti-TER119 vs. IgG control to induce anemia or control conditions, respectively. Anti-TER119 injections induced significant decreases in hemoglobin levels by 48 hours, as compared to control IgG injections (B). ICH was induced followed by serial MRI neuroimaging over 24 hours after ICH (C). Anti-TER119/anemic mice exhibited greater ICH lesion expansion as compared to controls (D). *: p<0.05; ****: p<0.001

**Supplemental table 1:**
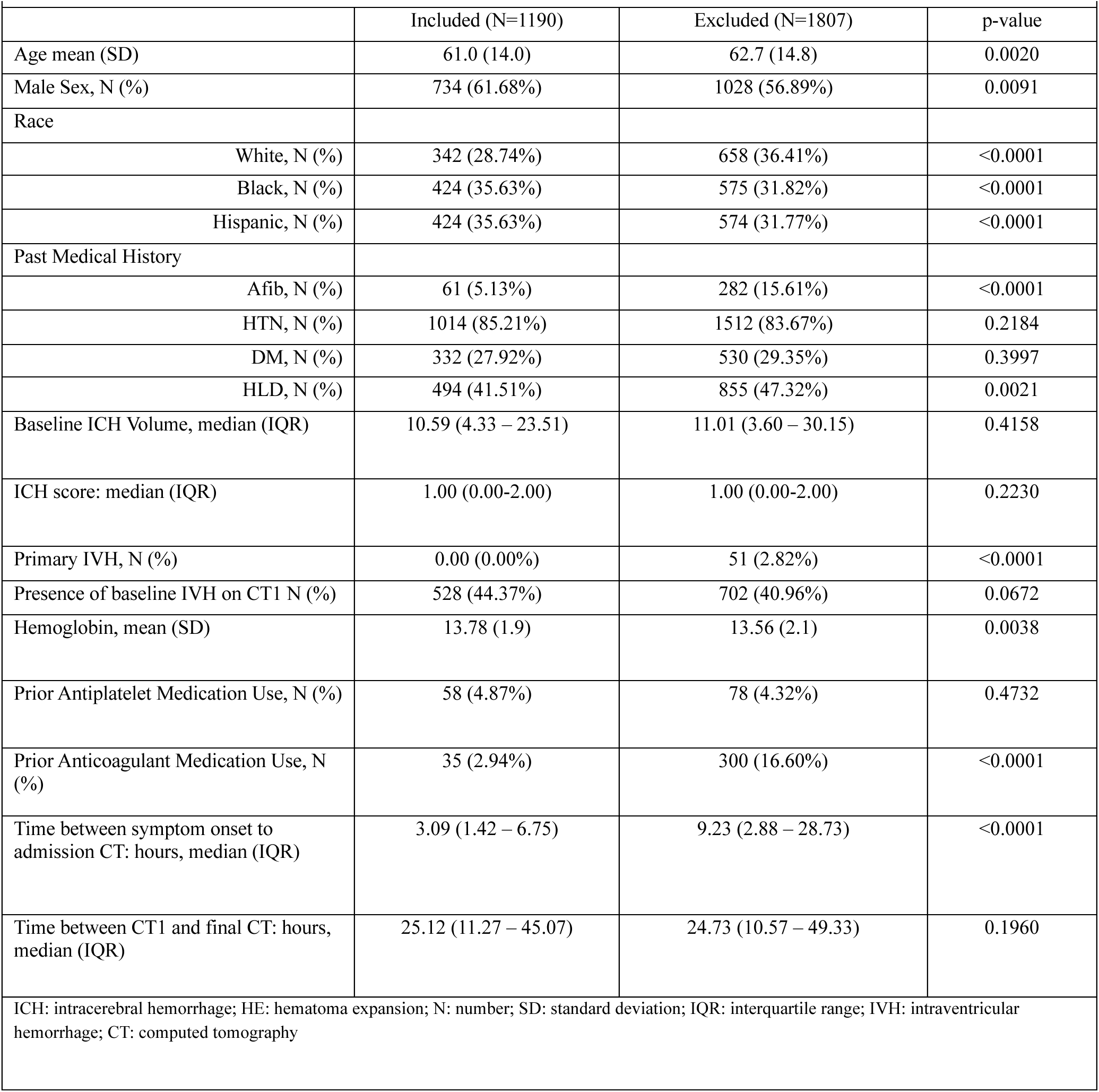
Intergroup differences between inclusion and exclusion cohort.

**Supplemental table 2:**
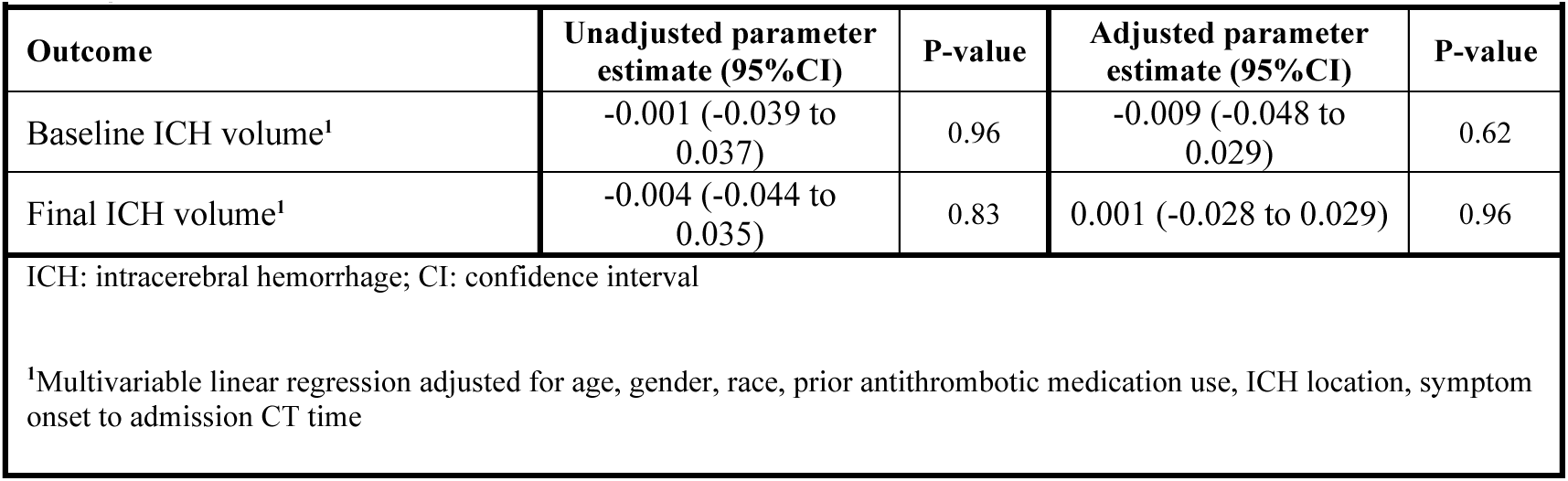
Crude and adjusted multivariable linear regression analysis assessing association of hemoglobin with ICH volume.

