## Supplementary figures and images for "Low hemoglobin causes hematoma expansion and poor intracerebral hemorrhage outcomes"

### Supplementary Figure 1

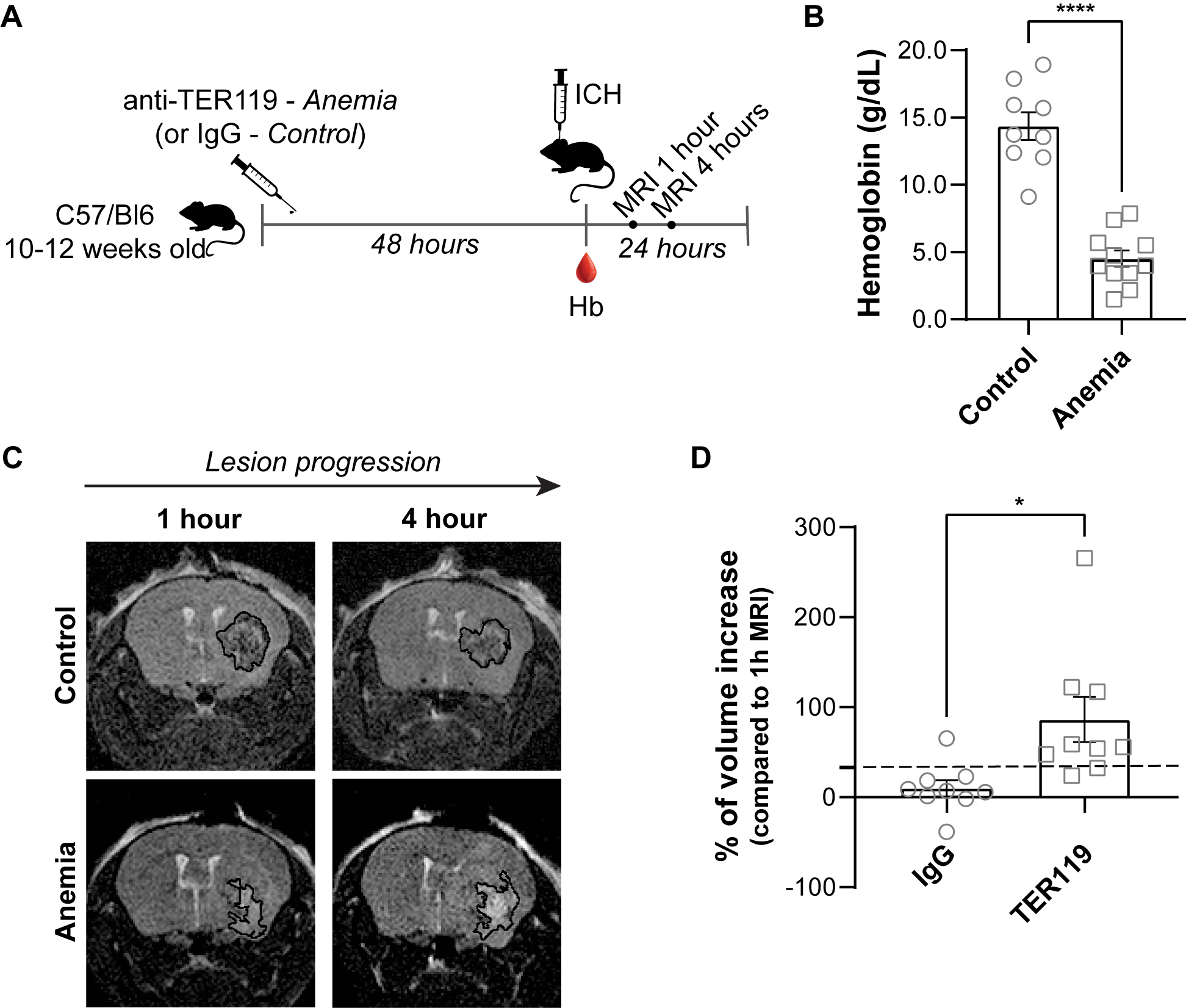
