## Supplementary Table 1 for "Low hemoglobin causes hematoma expansion and poor intracerebral hemorrhage outcomes"

| Supplemental table 1: Intergroup differences between inclusion and exclusion cohort | | | |
| --- | --- | --- | --- |
|  | Included (N=1190) | Excluded (N=1807) | p-value |
| Age mean (SD) | 61.0 (14.0) | 62.7 (14.8) | 0.0020 |
| Male Sex, N (%) | 734 (61.68%) | 1028 (56.89%) | 0.0091 |
| Race |  |  |  |
| White, N (%) | 342 (28.74%) | 658 (36.41%) | <0.0001 |
| Black, N (%) | 424 (35.63%) | 575 (31.82%) | <0.0001 |
| Hispanic, N (%) | 424 (35.63%) | 574 (31.77%) | <0.0001 |
| Past Medical History |  |  |  |
| Afib, N (%) | 61 (5.13%) | 282 (15.61%) | <0.0001 |
| HTN, N (%) | 1014 (85.21%) | 1512 (83.67%) | 0.2184 |
| DM, N (%) | 332 (27.92%) | 530 (29.35%) | 0.3997 |
| HLD, N (%) | 494 (41.51%) | 855 (47.32%) | 0.0021 |
| Baseline ICH Volume, median (IQR) | 10.59 (4.33 – 23.51) | 11.01 (3.60 – 30.15) | 0.4158 |
| ICH score: median (IQR) | 1.00 (0.00-2.00) | 1.00 (0.00-2.00) | 0.2230 |
| Primary IVH, N (%) | 0.00 (0.00%) | 51 (2.82%) | <0.0001 |
| Presence of baseline IVH on CT1 N (%) | 528 (44.37%) | 702 (40.96%) | 0.0672 |
| Hemoglobin, mean (SD) | 13.78 (1.9) | 13.56 (2.1) | 0.0038 |
| Prior Antiplatelet Medication Use, N (%) | 58 (4.87%) | 78 (4.32%) | 0.4732 |
| Prior Anticoagulant Medication Use, N (%) | 35 (2.94%) | 300 (16.60%) | <0.0001 |
| Time between symptom onset to admission CT: hours, median (IQR) | 3.09 (1.42 – 6.75) | 9.23 (2.88 – 28.73) | <0.0001 |
| Time between CT1 and final CT: hours, median (IQR) | 25.12 (11.27 – 45.07) | 24.73 (10.57 – 49.33) | 0.1960 |
| ICH: intracerebral hemorrhage; HE: hematoma expansion; N: number; SD: standard deviation; IQR: interquartile range; IVH: intraventricular hemorrhage; CT: computed tomography | | | |
