## Supplementary Table 2 for "Low hemoglobin causes hematoma expansion and poor intracerebral hemorrhage outcomes"

| **Supplemental Table 2:** Crude and adjusted multivariable linear regression analysis assessing association of hemoglobin with ICH volume | | | | |
| --- | --- | --- | --- | --- |
| **Outcome** | **Unadjusted parameter estimate (95%CI)** | **P-value** | **Adjusted parameter estimate (95%CI)** | **P-value** |
| Baseline ICH volume**^1^** | -0.001 (-0.039 to 0.037) | 0.96 | -0.009 (-0.048 to 0.029) | 0.62 |
| Final ICH volume**^1^** | -0.004 (-0.044 to 0.035) | 0.83 | 0.001 (-0.028 to 0.029) | 0.96 |
| ICH: intracerebral hemorrhage; CI: confidence interval  **^1^**Multivariable linear regression adjusted for age, gender, race, prior antithrombotic medication use, ICH location, symptom onset to admission CT time | | | | |
