## Supplementary material for "Low hemoglobin causes hematoma expansion and poor intracerebral hemorrhage outcomes": Table 1

| **Table 1:** Baseline Characteristics of ICH patients with and without HE | | | | |
| --- | --- | --- | --- | --- |
|  | **All ICH**  **N=1190** | **HE**  **N=244** | **No HE**  **N=946** | **P-value** |
| **Age:** mean (SD) | 61 (14) | 62 (14) | 61 (14) | 0.25 |
| **Male:** N (%) | 734 (62) | 167 (68) | 567 (60) | 0.01 |
| **Race:** N (%) |  |  |  | 0.02 |
| White | 343 (29) | 85 (35) | 258 (27) |  |
| Black | 424 (36) | 70 (29) | 354 (37) |  |
| Hispanic | 423 (36) | 89 (36) | 334 (35) |  |
| **Medical History:** N (%) |  |  |  |  |
| Dyslipidemia | 494 (42) | 105 (43) | 389 (41) | 0.69 |
| Atrial Fibrillation | 61 (5) | 21 (9) | 40 (4) | 0.01 |
| Hypertension | 1013 (85) | 204 (84) | 809 (86) | 0.45 |
| Diabetes | 332 (28) | 64 (26) | 268 (28) | 0.51 |
| **Medication History:** N (%) |  |  |  |  |
| Antiplatelet | 58 (5) | 18 (7) | 40 (4) | 0.04 |
| Anticoagulant | 35 (3) | 8 (3) | 27 (3) | 0.73 |
| **Clinical/Radiographic** |  |  |  |  |
| ICH Score: median (IQR) | 1 (0-2) | 1 (0-2) | 1 (0-2) | 0.99 |
| Baseline ICH volume (mL): median (IQR) | 10.6 (4.3-23.5) | 12.4 (4.2-28.9) | 10.1 (4.4-22.1) | 0.07 |
| IVH presence: N (%) | 528 (44) | 82 (34) | 446 (47) | <0.001 |
| Time to admission CT (hours): median (IQR)** | 3.1 (1.4-6.8) | 1.9 (1.2-4.8) | 3.6 (1.6-7.4) | <0.001 |
| Time between baseline and final CT (hours): median (IQR) | 25.1 (11.3-45.1) | 24.8 (12.9-42.0) | 25.3 (11.1-45.4) | 0.98 |
| **Hemoglobin** (g/dL): mean (SD) | 13.8 (1.9) | 13.6 (2.0) | 13.8 (1.9) | 0.14 |
| ICH: intracerebral hemorrhage; HE: hematoma expansion; N: number; SD: standard deviation; IQR: interquartile range; IVH: intraventricular hemorrhage; CT: computed tomography | | | | |
